# Cultured *Ostreococcus* is cobalamin-independent

**DOI:** 10.1101/065771

**Authors:** Mark Paul Pašek

## Abstract

The marine microalga *Ostreococcus* is considered to depend on the methionine synthase METH and its methylated cobalamin cofactor for methionine synthesis because it lacks METE, the methylcobalamin-independent methionine synthase. Here I describe minimal media lacking both biotin and cobalamin yet suitable for growth of cultured *Ostreococcus*.

## Introduction

The marine unicellular alga *Ostreococcus* has evolved clades A, B, and C, each with sequenced representatives, as well as clade D, *Ostreococcus mediterraneus* (Subirana et al., 2013). The model of methionine biosynthesis in *Ostreococcus* states that METH [methylcobalamin-dependent] is the sole methionine synthase and is based on the identification of the gene encoding METH in all three clades. In addition, the gene encoding METE [5-methyltetrahydrofolate-dependent; cobalamin-independent] appears missing for all three clades, leading to the prediction of cobalamin-dependent growth (Helliwell et al., 2011). These observations suggest that if *Ostreococcus* cells were transformed with synthetic DNA designed to express METE, transformants would utilize the endogenous METH substrate 5-methyltetrahydofolate to synthesize methionine and could be selected in minimal medium in the absence of cobalamin.

## Materials and Methods

*Ostreococcus* cell cultures were obtained from the Roscoff Culture Collection, Roscoff, France, and the Provasoli-Guillard National Center for Marine Algae and Microbiota, East Boothbay, Maine, USA. Panels of candidates to be evaluated for DNA transformation were assembled. The panel for clade A consisted of NCMA2972 (Palenik et al., 2007), RCC754, RCC755, and RCC756. The panel for clade C included RCC614 and RCC745 (Derelle et al., 2006). The members of the panel for clade D were RCC789, RCC1107, RCC1119, and RCC1121.

*Ostreococcus* cells were cultured in suspension by rocking under continuous blue illumination in minimal medium at room temperature. Minimal medium is artificial seawater (McDonald et al., 2010) supplemented with 0.9 mM NaNO_3_, 100 µM Na_2_EDTA•2H_2_O, 10 µM NaH_2_PO_4_•H_2_O, 10 µM FeNaEDTA, 0.9 µM MnCl_2_•4H_2_O, 300 nM thiamine•HCl, 100 nM Na_2_SeO_3_, 80 nM ZnSO_4_•7H_2_O, 30 nM Na_2_MoO_4_•2H_2_O, and 10 nM CuSO_4_•5H_2_O. This list of supplements is simplified from that formulated by Keller (Keller et al., 1987), most notably lacking biotin, cyanocobalamin, CoSO_4_•7H_2_O, NH_4_Cl, and tris(hydroxymethyl)aminomethane. In addition, the selenite concentration is 10 nM in Keller medium.

For polymerase chain reaction DNA amplification, template DNA was prepared with Quick-gDNA^™^ from Zymo Research, PCR primer pairs were synthesized by Integrated DNA Technologies, and targets were amplified with the Q5 DNA polymerase from New England BioLabs.

## Results

Using DNA from RCC614 and RCC745 as template and two primer pairs for each of the two targets *METH* and *URA5/3*, polymerase chain reaction DNA amplification yielded in each case DNA fragments of the predicted size, and authenticated cell culture identity as *Ostreococcus* after long-term serial culture in minimal medium with 400 pM cobalamin. For NCMA2972, a primer pair also targeting *URA5/3* was used (Table 1).

**Table 1.**
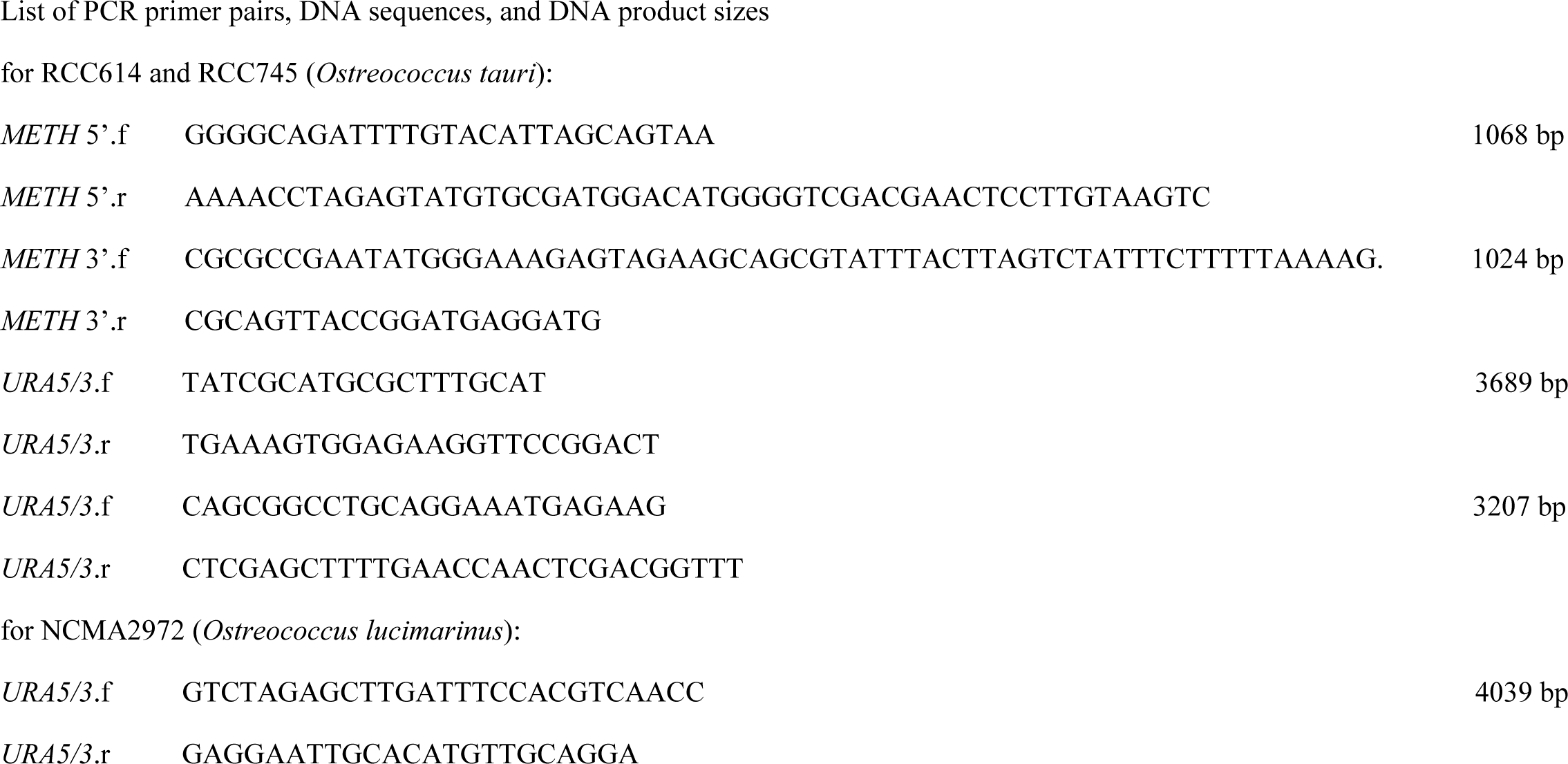
List of PCR primer pairs, DNA sequences, and DNA product sizes for RCC614 and RCC745 (*Ostreococcus tauri*):

Minimal medium with cobalamin supported extended serial growth for each cell culture in each clade. In addition, minimal medium containing cobalamin and solidified with 0.125% gellan gum (Sigma-Aldrich) supported diverse green bacterioplanktonic surface growth (Figure 1) for each cell culture in each clade, with the exception of NCMA2972.

**Figure 1.**
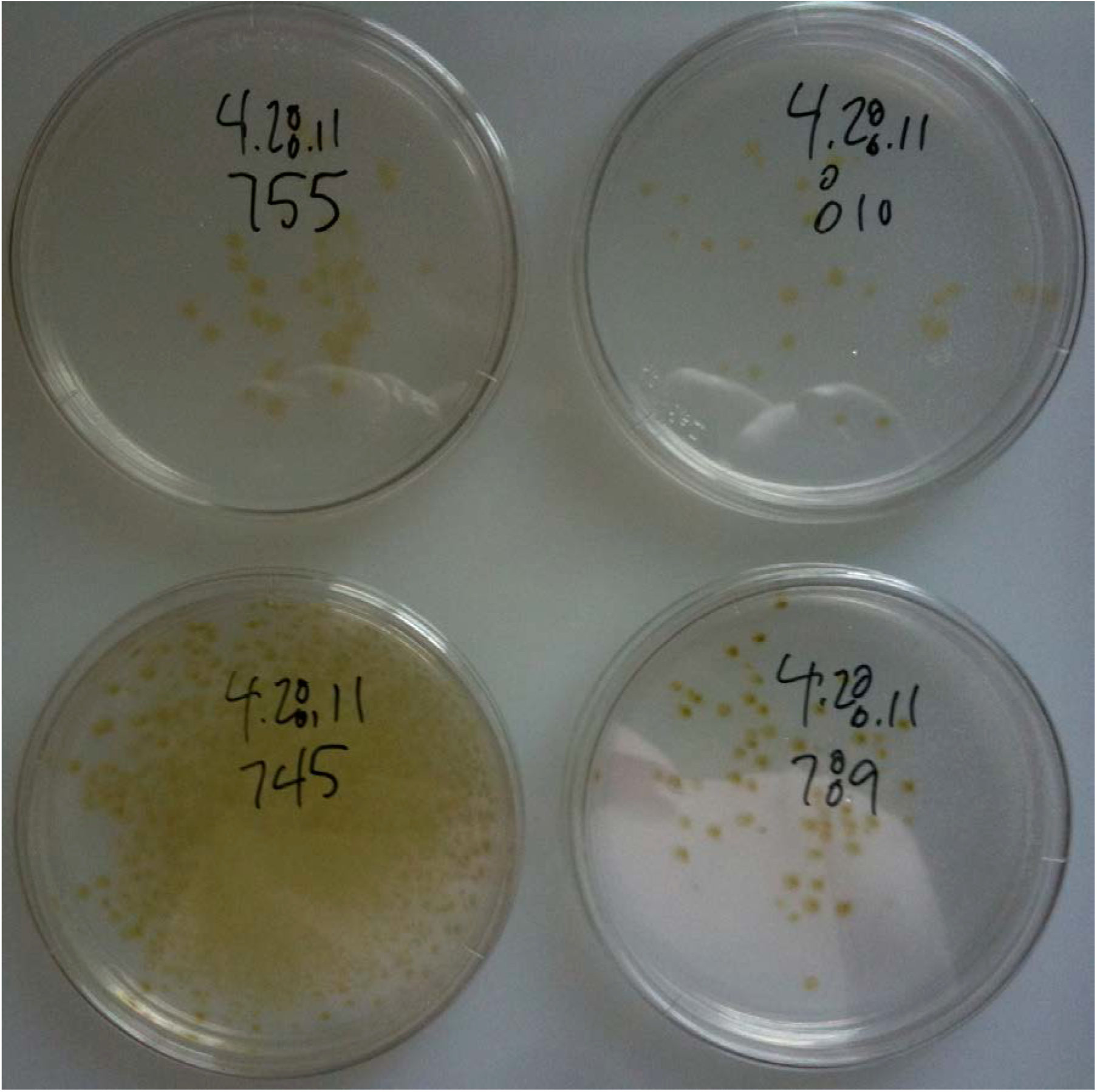
Bacterioplanktonic growth of *Ostreococcus* clades A, C, and D on minimal medium with cobalamin solidified with 0.125% gellan gum.

As a transformation host, RCC614 was chosen. Mock transformation experiments, without synthetic DNA, were designed to quantitate and evaluate background growth on solidified minimal medium lacking cobalamin. RCC614 suspension cells were cultured in minimal medium with 400 pM cobalamin. 10^6^ cells (100 µl) were mixed with 50 µl CutSmart^®^ restriction enzyme buffer [50 mM potassium acetate-10 mM magnesium acetate-20 mM tris(hydroxymethyl)aminomethane acetate (pH 7.9)-100 µg/ml bovine serum albumin; New England BioLabs], layered onto 100 µl protoplast transformation buffer [100 mM CaCl_2_-200 mM mannitol-40% PEG4000; Yoo et al., 2007], briefly vortexed, incubated for 5 minutes at room temperature, diluted into 10 ml of minimal medium containing 20 pM cobalamin, and cultured for 24 hours. Cells were then aerosolized onto minimal media lacking cobalamin solidified with 0.125% gellan gum at a density of 10^5^ cells per 90 mm plate (20 ml solid media per plate). After 10 days at room temperature, approximately 10 to 20 green bacterioplanktonic colonies per plate emerged at normal growth rates.

Twelve cobalamin-independent colonies were picked into minimal medium without cobalamin and cultured; all cultured remained cobalamin-independent after more than 10 serial passages (1/200 dilution/passage). Each cobalamin-independent culture was authenticated as RCC614 by polymerase chain reaction DNA amplification using the primer pair for *URA5/3* listed in Table 1 which yields a 3207 bp diagnostic fragment (Figure 2).

**Figure 2.**
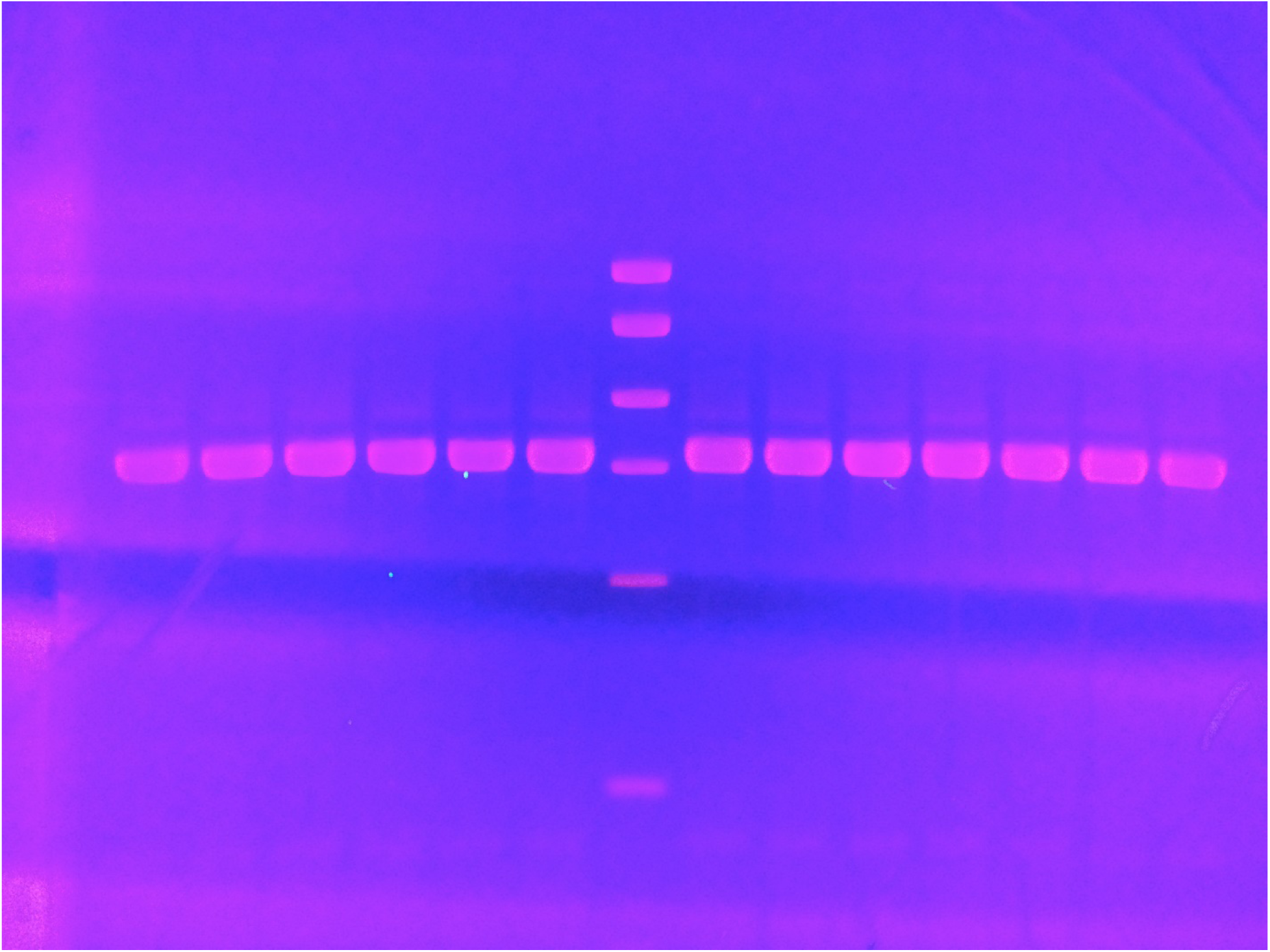
PCR authentication of cobalamin-independent clones as *Ostreococcus tauri* RCC614. Lanes 1-6 and 9-14 correspond to cobalamin-independent clones isolated on gellan gum-solidified minimal medium lacking cobalamin. Lane 8 is RCC614. Lane 7 is the Mass DNA ladder from New England BioLabs. The diagnostic 3207 bp DNA fragment is synthesized with a primer pair targeting the *URA5/3* locus (Table 1).

It is concluded that cobalamin-independent *Ostreococcus* colonies are obtained at a frequency of approximately 0.01% when selected on gellan gum-solidified minimal medium without cobalamin.

## Discussion

The utility of *METE* as a positive selection marker relies on a cobalamin-dependent *Ostreococcus* transformation recipient. RCC614 is a representative co-culture of *Ostreococcus* and various phycosphere-associated marine bacteria (Abby et al., 2014; Lupette et al., 2016). The bacteria may supply biotin, cobalamin, and methionine to the *Ostreococcus* phycosphere. A mock transformation as described above of RCC614 will yield *Ostreococcus* bacterioplanktonic colony growth on minimal medium with no added biotin and cobalamin, as observed. Only a cobalamin-dependent *Ostreococcus* cell culture, not necessarily biotin-independent, is required to implement the use of *METE* as a positive selection marker.

Both RCC614 and *Ostreococcus* RCC809 (Grigoriev, 2011; clade B) lack the gene encoding the algal-specific cobalamin acquisition protein 1, whereas NCMA2972 has it (Bertrand et al., 2012). Perhaps *Ostreococcus* clades C and B obtain methionine from their phycosphere while clade A can utilize cobalamin in addition.

